# Long-term effects of THC exposure on reward learning and motivated behavior in adolescent and adult male rats

**DOI:** 10.1101/2022.12.05.519170

**Authors:** Briac Halbout, Collin Hutson, Leann Hua, Victoria Inshishian, Stephen V. Mahler, Sean B. Ostlund

**Affiliations:** Department of Anesthesiology and Perioperative Care, School of Medicine, University of California, Irvine, Irvine, CA, 92697, USA; Department of Neurobiology and Behavior, School of Biological Sciences, University of California, Irvine, Irvine, CA, 92697, USA

**Keywords:** cannabis, marijuana, addiction, habit, goal-directed action, motivation, emotion, eating

## Abstract

**Rationale:** The endocannabinoid system makes critical contributions to reward processing, motivation, and behavioral control. Repeated exposure to THC or other cannabinoid drugs can cause persistent adaptions in the endocannabinoid system and associated neural circuitry. It remains unclear how such treatments affect the way rewards are processed and pursued.

**Objective and methods:** We examined if repeated THC exposure (5 mg/kg/day for 14 days) during adolescence or adulthood led to long-term changes in rats’ capacity to flexibly encode and use action-outcome associations for goal-directed decision making. Effects on hedonic feeding and progressive ratio responding were also assessed.

**Results:** THC exposure had no effect on rats’ ability to flexibly select actions following reward devaluation. However, instrumental contingency degradation learning, which involves avoiding an action that is unnecessary for reward delivery, was augmented in rats with a history of adult but not adolescent THC exposure. THC-exposed rats also displayed more vigorous instrumental behavior in this study, suggesting a motivational enhancement. A separate experiment found that while THC exposure had no effect on hedonic feeding behavior, it increased rats’ willingness to work for food on a progressive ratio schedule, an effect that was more pronounced when THC was administered to adults. Adolescent and adult THC exposure had opposing effects on the CB1-receptor dependence of progressive ratio performance, decreasing and increasing sensitivity to rimonabant-induced behavioral suppression, respectively.

**Conclusions:** Our findings reveal that exposure to a translationally relevant THC exposure regimen induces long-lasting, age-dependent alterations in cognitive and motivational processes that regulate the pursuit of rewards.

## Introduction

Cannabis is widely viewed as being innocuous despite negatively impacting the lives of many of its users. Nearly a third of regular cannabis users meet the diagnostic criteria for cannabis use disorder (Hasin et al. 2015), indicating a pattern of intake that persists in spite of its adverse consequences. This is on par with estimates of heroin and cocaine use disorder among regular users of these drugs (Ferland and Hurd 2020), which is worrisome given the high and growing prevalence of regular cannabis use (Hammond et al. 2021; Yu et al. 2020). Individuals with cannabis use disorder also face a high risk of relapse when attempting to quit (Copeland et al. 2001; Moore and Budney 2003), as with other addictive drugs. Such findings highlight the need to advance understanding of how cannabis comes to exerts such powerful control over behavior.

With repeated exposure, addictive drugs share a tendency to induce long-lasting behavioral and neural adaptations, and cannabis is no exception to this rule. Rats given repeated exposure to Δ^9^-tetrahydrocannabinol (THC), the main psychoactive component of cannabis, exhibit sensitization to the acute behavioral effects of this drug, as well as its ability to stimulate mesolimbic dopamine release (Cadoni et al. 2008). Some have argued that such adaptations reflect a fundamental change in motivational processing – termed incentive-sensitization – which fuels the addiction process by amplifying the desire to pursue drugs and other rewards (Berridge and Robinson 2016; Robinson and Berridge 1993).

An alternative view assumes that problematic, uncontrolled drug use reflects an underlying failure of adaptive, goal-directed decision making, which may be caused or exacerbated by repeated drug intake (Belin et al. 2013; Everitt and Robbins 2005; Ostlund and Balleine 2008). In line with this view, previous studies have shown that repeated exposure to various addictive drugs (Corbit et al. 2014; LeBlanc et al. 2013; Nelson and Killcross 2006; Nelson and Killcross 2013; Nordquist et al. 2007a; Renteria et al. 2018) including THC (Nazzaro et al. 2012) can promote the development inflexible reward-seeking habits (but see Ferland et al. 2022). Importantly, such effects are believed to reflect an acceleration in adaptive habit learning and need not involve a more wide-ranging impairment in goal-directed decision making or control (Ostlund and Balleine 2008). While the impact of repeated cannabinoid exposure on goal-directed control remains unclear, such treatments have been shown to cause long-lasting alterations in the structure and function of the prefrontal cortex (Cass et al. 2014; Renard et al. 2016; Rubino et al. 2015), a critical hub for goal-directed behavior (Balleine 2019; Bradfield and Hart 2020; Turner and Parkes 2020).

The endocannabinoid system undergoes profound changes during adolescence which are believed to help shape normal neurocognitive development but also make the brain more vulnerable to the harmful effects of cannabis and other drugs (Crews et al. 2007; Schneider 2008; Spear 2016; Stringfield and Torregrossa 2021a). Heavy adolescent cannabis use has been linked to later cognitive impairment and mental health problems (Chadwick et al. 2013; Levine et al. 2017; Lubman et al. 2015; Volkow et al. 2016). Likewise, animal studies have shown that adolescent exposure to cannabinoid drugs can cause persistent alterations in behavioral measures of emotion, motivation, and cognition (Bambico et al. 2010; Cha et al. 2007; Gleason et al. 2012; Higuera-Matas et al. 2015; Jacobs-Brichford et al. 2019; Kruse et al. 2019; O’Shea et al. 2004; Quinn et al. 2008b; Realini et al. 2011; Renard et al. 2017b; Rubino et al. 2008; Scherma et al. 2016; Schneider and Koch 2003; Schoch et al. 2018; Zamberletti et al. 2012), often having more pronounced effects than adult cannabinoid exposure (Quinn et al. 2008b; Schneider and Koch 2003).

Research on the effects of cannabinoid exposure on reward-related behavior have produced mixed results that appear to depend on features of the drug exposure regimen (Stringfield and Torregrossa 2021a). For instance, long-term suppression of reward consumption and reward-motivated behavior has been observed following adolescent exposure to potent synthetic cannabinoids (Bambico et al. 2010) or high doses of THC (e.g., twice-daily injections of up to 10 mg/kg)(Realini et al. 2011; Rubino et al. 2008; Scherma et al. 2016). Such findings are notable because anhedonia – i.e., diminished interest in healthy, reward-seeking activities – is a common feature of depression, anxiety, and other psychiatric disorders (Husain and Roiser 2018). In contrast, repeated adolescent exposure to lower doses of THC (2.5-5 mg/kg/day) have often been seen to elevate, rather than suppress, reward-motivated behavior (Kruse et al. 2019; Orihuel et al. 2021). Further research on the effects of more moderate THC dosing regimens is therefore warranted, particularly given their relevance to modeling early-stage, cannabis use in humans (Poulia et al. 2021; Ruiz et al. 2021; Torrens et al. 2020).

The current study investigated the long-term consequences of an intermediate-dose THC regimen (5 mg/kg/day for 14 days) on reward-related behavior in male rats, using selective assays of emotion (hedonic feeding), motivation (effort exertion), and behavioral control (goal-directed decision making). Both adolescent- and adult-THC exposure conditions were included to investigate whether adolescent development exacerbates or otherwise alters the long-term behavioral effects of this treatment.

We also conducted a dose-response analysis of the behavioral effects of the inverse CB1-receptor agonist rimonabant on reward-motivated behavior in rats with or without a history of repeated THC exposure. Adolescent THC exposure is known to decrease expression and function of the endocannabinoid CB1 receptor (Kruse et al. 2019; Rubino et al. 2015; Silva et al. 2016; Stringfield and Torregrossa 2021b; Zamberletti et al. 2012), which is critical for reward-motivated behavior (Cha et al. 2007; Friemel et al. 2014; Hernandez and Cheer 2012; Maccioni et al. 2008; Marusich and Wiley 2012; Rasmussen and Huskinson 2008; Solinas and Goldberg 2005; Ward and Dykstra 2005). We therefore hypothesized that THC exposure, particularly during adolescence, would alter the CB1 receptor-dependence of reward-motivated behavior.

## Material & Methods

All procedures were approved by the UC Irvine Institutional Animal Care and Use Committee (IACUC) and were in carried out in accordance with the National Research Council Guide for the Care and Use of Laboratory Animals.

### Animals

Male Long-Evans rats (N = 80) were obtained from Charles River. For adolescent exposure cohorts, rats were weaned at postnatal day (PD) 21 and arrived at our facility at PD 22. Adult exposure cohorts arrived aged approximately 12 weeks. Adult rats were pair-housed throughout the study, whereas adolescent rats were initially housed in groups of four before being pair-housed one week prior to behavioral testing (see below for detailed experimental designs). Rats were housed in transparent plastic cages in a temperature- and humidity-controlled vivarium. The rats were tested during the light phase of a standard 12:12 h light:dark schedule, and had ad libitum access to food and water in their home cages throughout the experiment, except when food restricted for specific procedures as indicated below.

### Apparatus

Behavioral procedures were conducted in identical operant chambers (ENV-007, Med Associates, St Albans, VT, USA), each housed in a sound- and light-attenuated cubicle. A food-delivery port was located at the center of one end-wall of the chamber, 2.5 cm above the stainless-steel grid floor. Separate cups within the food port were used to deliver sweetened condensed milk (SCM) solution via a syringe pump located outside of the cubicle or 45-mg grain pellets (BioServ) via an automated pellet dispenser. A photobeam detector positioned across the food-port entrance was used to monitor head entries. SCM licking responses were continuously recorded during consumption test sessions using a contact lickometer device (ENV-250B, Med Associates, St Albans, VT, USA).

Locomotor activity was monitored with four photobeams that were positioned in a horizontal plane ∼2 cm above the grid floor. Each chamber was also equipped with two retractable levers positioned to the left and right of the food port. A houselight (3 W, 24 V) at the top of the opposite end-wall provided general illumination and a fan mounted on the cubicle provided ventilation and background noise. Experimental events were controlled and recorded with a 10-msec resolution using MED-PC IV software.

### Drug preparation and treatment

THC was provided by the NIDA Drug Supply Program and was prepared daily by evaporating vehicle under N_2_ and dissolving to dose in 5% Tween 80 in saline (1 ml/kg) prior to intraperitoneal injections (Burston et al. 2010; Ruiz et al. 2021; Torrens et al. 2020). Rats were administered a series of 14 once-daily i.p. injections of THC or vehicle beginning at PD 30 for adolescent exposure cohorts (THC n=20; Veh n=20), or at 13 weeks postnatal for adult exposure cohorts (THC n=20; Veh n=20). Rimonabant (ApexBio Technology) was prepared fresh each day, dissolved in Tween 80/PEG-400/sterile 0.9% saline (1:1:18, vol/vol/vol) and sonicated for 10 min at 30°C. Vehicle and rimonabant suspension were injected at a volume of 1 mL/kg.

### Experiment 1

#### Overview

Rats (N = 40) were pretreated with THC or vehicle as adolescents or adults, using a fully factorial design (n’s = 10/group). This experiment was run in two replications with fully balanced groups. Behavioral testing began after a washout period of 63-67 days. Rats were food restricted for 3 d before and throughout testing by providing each animal with 10-14 g of home chow at the end of each day to maintain them at ∼85% of their estimated free-feeding bodyweight. We then investigated the effects of THC pre-exposure on the acquisition and control of instrumental reward-seeking behavior. We specifically probed rats’ ability to make flexible choices between actions based on changes in reward value and action-outcome contingency, which are hallmarks of goal-directed control (Balleine and Dickinson 1998).

#### Instrumental action-outcome training

Rats initially received 2 days of magazine training, during which 20 grain pellets and 20 deliveries of 120 uL of 50% SCM were delivered on a random time (RT) 30-s schedule with the levers retracted. Rats were then given 10 days of instrumental training on two distinct action–outcome contingencies (i.e., R1 → O1 and R2 → O2). The left and right lever-press actions were trained in separate sessions each day.

Lever-outcome arrangements were counterbalanced with drug treatment conditions, such that, for half of the rats in each group, pressing the left lever produced SCM solution and pressing the right lever produced grain pellets, whereas the other half received the opposite arrangement. Only the active lever (left or right) was extended during individual training sessions, which terminated after 30 min elapsed or 20 rewards were earned. Rats were placed in their home cage for at least 2 h between the two daily training sessions. The schedule used to reinforce lever pressing began with 2 days of fixed ratio (FR)-1 training, but then shifted through a series of increasingly more effortful random ratio (RR) schedules, with 2-day intervals of RR-5, RR-10, and RR-20 training, such that an average of 20 presses were needed to earn each reward during the last phase of instrumental training. Previous studies have shown that rats trained with similar protocols involving multiple action-outcome contingencies tend to prevent habit formation even after overtraining (Colwill and Rescorla 1985; Colwill and Triola 2002; Halbout et al. 2016; Kosaki and Dickinson 2010).

#### Reward devaluation

We used a specific-satiety procedure to assess the effects of post-training reward devaluation on rats’ choice between the two instrumental reward-seeking actions, as in our previous publications (Halbout et al. 2016; Halbout et al. 2019; Kosheleff et al. 2018). On test days, rats were given 60 min of unrestricted access to 50% SCM or grain pellets (counterbalanced with pretreatment groups) in their home cages. Rats were then placed in the behavioral chambers for a 15-min test session, during which they had continuous access to both levers. Each test began with a 5-min extinction phase, such that lever presses were recorded but were not reinforced, which was done to probe action selection in the absence of explicit response-contingent feedback, thereby requiring retrieval of previously encoded action-outcome associations (Balleine and Dickinson 1998). This was immediately followed by a 15-min rewarded phase, during which each action was reinforced with its respective outcome. An FR-1 schedule was in place for the first 5 reward deliveries before shifting to a RR-20 schedule for the remainder of the session. The rewarded phase was included as a tool to confirm the efficacy of the specific satiety procedure and to determine rats’ ability to select actions when provided with explicit feedback about the consequences of their actions, which can further promote goal-directed control (Ostlund and Balleine 2008). After the first test, rats were given a session of retraining with each lever using a schedule of reinforcement that shifted within-session with reward delivery, moving from FR-1 (first 3 rewards) to RR-5 (next 5 rewards) to RR-10 (1 reward), before reaching the RR-20 schedule that would remain in place for the remainder of the session. Retraining sessions lasted for 30 min or until 20 rewards were earned. On the following day, rats were given a second reward devaluation test after being satiated on the alternative food outcome.

#### Action-outcome reversal training and reward devaluation

Following the first round of devaluation testing, rats were given 5 days of instrumental retraining using a RR-20 reinforcement schedule. These sessions used the same parameters as initial instrumental training sessions (see above), except that the original action-outcome contingencies were reversed (i.e., R1 → O2 and R2 → O1), such that the lever that had once produce SCM now produced grain pellets, and vice versa. Rats were then administered a second round of two reward devaluation tests (one with SCM devalued and one with pellets devalued; counterbalanced), as described above, with a session of retraining in between tests using the reversed contingencies. This post-reversal test assays rats’ capacity to use recency to disambiguate conflicting action-outcome associations for flexible, goal-directed decision making (Bradfield and Balleine 2017; Panayi and Killcross 2018; Parkes et al. 2018).

#### Contingency degradation training and testing

Rats were then retrained on the original action-outcome contingencies (i.e., R1 → O1 and R2 → O2) on an RR-20 schedule for 3 days, as described above. Next, they were given 10 days of instrumental contingency degradation training (Balleine and Dickinson 1998; Corbit et al. 2002; Halbout et al. 2016), which was used to selectively weaken the predictive relationship for one of the two action-outcome contingencies while continuing to reinforce both actions with their original outcomes on a modified RR-20 schedule. Specifically, sessions were divided into a series of 1-s periods and the first press performed in each period had a 1-in-20 chance of producing reward [p(outcome/response) = 0.05]. As before, the two actions were trained in separate daily sessions, though these sessions were now limited to 20 min without a cap on the number of rewards that could be earned. Importantly, one of the two outcomes (SCM or grain pellets) was also delivered in a noncontingent manner. Specifically, the noncontingent outcome had a 1-in-20 chance of being delivered at the end of any 1-s period without a lever-press response [p(outcome/no response) = 0.05]. The identity of the noncontingent outcome was fixed for individual subjects and was counterbalanced across groups. This outcome was noncontingently delivered in all contingency degradation training sessions, regardless of which lever was available, such that, for degraded sessions, the noncontingent outcome was the same as the response-contingent outcome [e.g., R1 → O1 & No R1 → O1], whereas for non-degraded sessions, the noncontingent outcome was different from the response-contingent outcome [e.g., R2 → O2 & No R2 → O1]. Thus, one action (R1) lost its predictive value because its outcome was just as likely to occur after that action as in its absence, whereas the alternative action remained a unique and reliable predictor of its outcome. Five-min extinction tests were conducted after the 6^th^ and 10^th^ day of contingency degradation training to assess how this procedure altered rats’ choice between actions. At test, both levers were continuously available in the absence of either response-contingent or noncontingent reward delivery.

### Experiment 2

#### Overview

A separate set of rats (N = 40) were pretreated with THC or vehicle as adolescents or adults using a fully factorial design (n’s = 10/group). This experiment was run in two replications with fully balanced groups. Behavioral testing began after a washout period of 23-28 days. Thereafter, we assessed the effects of THC pre-exposure on the hedonic component of feeding during sessions of free access to varying concentrations of SCM solution. We then assessed their motivation, or willingness to exert effort, for SCM using an operant progressive ratio task. After establishing baseline performance, we conducted a dose-response analysis of the response-suppressive effects of rimonabant (CB1 receptor inverse agonist) to assess alterations in CB1 receptor-dependent motivational function. Behavioral testing in this experiment was conducted without food or water restriction except as described below.

#### Hedonic feeding

Rats were handled for 3 days prior to testing. During the last 2 days of handling, rats were also given 2h of free access to a bottle containing 50% SCM to familiarize them with this solution. They were then given 4 daily 30-min sessions to provide them with experience consuming 50% SCM in the behavior chambers. At the beginning of each session, the food cup was filled with 30 ul of SCM over a 0.5-sec interval via syringe-pump activation. Cups were refilled whenever rats drank the solution. Specifically, any lick response detected when the syringe pump was inactive resulted in the immediate injection of a new 15-ul volume of SCM, delivered over 0.25 sec. Licks detected when the syringe pump was already active were recorded but did not influence the ongoing SCM delivery. This contingency ensured that rats had continuous access to SCM at a maximal delivery rate of 60 ul/sec while preventing the cup from being overfilled.

To assess the influence of reward palatability on feeding behavior, rats were given a series of 8 separate 90-min sessions of access to varying concentrations of SCM (5, 10, 25 and 50%). Test order was pseudorandom (latin square) and counterbalanced. Our primary measure of hedonic feeding was bodyweight-normalized SCM intake (ml/kg) during the first 3 min of active licking behavior (beginning after the first contact with SCM), as in our previous publications (Marshall et al. 2017). This and related measures selectively track the influence of taste palatability on fluid intake while avoiding the inhibitory effects of post-ingestive satiety (Davis and Perez 1993; Davis and Smith 1988). Rates of licking (licks/min) and locomotor behavior (breaks/min) during tests sessions were also analyzed.

#### Progressive ratio

After assessing hedonic feeding behavior, rats were given instrumental training to lever press for 50% SCM reward (120 ul). Each session ended after 30 min or 20 rewards were earned. Rats were reinforced on a FR-1 schedule during the first 2 sessions and an FR-3 schedule during the next 2 sessions. To facilitate acquisition of the lever-press response, rats were mildly food restricted during these initial FR training sessions. Home chow was removed from their cages the night before the first session and rats were given 2 h of access to chow per day after each training session. Unrestricted access to chow in the home cage was resumed after the last FR-3 session.

Rats were then trained on a progressive ratio (PR)-3 schedule of reinforcement, with a response requirement that began at 1 press and progressed in 3-press increments for each reward earned in that session (i.e., 1, 4, 7, etc.). Rats received 6 initial days of PR-3 training sessions, each lasting 90 min with no limit on the number of rewards that could be earned. The concentration of SCM was set at 50% for all but the 5^th^ session, when it was shifted to 5% to assess the effect of reward palatability on task performance.

#### Effects of rimonabant on progressive ratio performance

We then assessed the effects of disrupting CB1 receptor activity on PR-3 task performance, using the same testing procedures described above, with a 50% SCM reward. Thirty-min before each test session, rats were pretreated with varying doses of rimonabant (0, 0.3, 1 and 3 mg/kg, i.p.). Each test was followed by at least 1 day off to allow for drug washout and was preceded by 1 day of drug-free retraining on the PR-3 task.

### Data analysis

Data were analyzed with univariate or repeated measures ANOVAs in SPSS v. 28. Significance was set at p < .05. Significant interactions were followed by an analysis of lower-order interactions or simple effects, as appropriate, to identify contributing factors. Bodyweights on the first and last day of drug exposure and at the beginning of behavioral testing (following drug washout) were analyzed using a two-way ANOVA with Drug (THC vs. vehicle exposure) and Age (adolescent vs. adult) as factors. Instrumental response rates (presses per minute) were analyzed using mixed ANOVAs which included Drug and Age as factors in addition to other within-subjects factors as appropriate. Initial Action-Outcome data were averaged across levers and sessions within each phase of reinforcement before they were analyzed with a Drug x Age x Schedule ANOVA. Subsequent retraining data were analyzed with Drug x Age x Session ANOVAs. Reward devaluation test data were averaged across test-pairs (one with each reward devalued) and analyzed separately for each test phase (extinction vs. reinforced) using Drug x Age x Devaluation (devalued vs. nondevalued action) ANOVAs. Instrumental contingency degradation data were normalized to baseline response rates (average of last 3 days of retraining) and were analyzed using a Drug x Age x Degradation (degraded vs. nondegraded action) x Session (1-12) ANOVA. Equipment malfunction during contingency degradation training led to a modest loss of data (20 of 800 total data points, or 2.5%), which was dispersed across groups. Missing values were interpolated for data analysis by taking the average response rate in surrounding sessions. Data from the first and second degradation test were analyzed separately using Drug x Age x Degradation ANOVAs. For hedonic feeding tests, we analyzed the temporal pattern of SCM intake (bodyweight normalized) using a Drug x Age x Time (1-30 mins) ANOVA. Early (first 3 min) and total intake were also analyzed using a Drug x Age x Concentration (5%, 10%, 25%, 50%) ANOVA, as was total locomotor activity.

Number of rewards earned served as our main measure of progressive ratio performance. The effect of SCM concentration on progressive ratio performance was analyzed using a Drug x Age x Concentration (5% vs. 50%) ANOVA. The effect of rimonabant pretreatment on progressive ratio performance was analyzed using a Drug x Age x Dose (0, 0.3, 1, 3 mg/kg) ANOVA, which was followed by an orthogonal polynomial trend analysis to characterize linear and nonlinear dose-response functions (Randall et al. 2011; Wickens and Keppel 2004).

## RESULTS

### Experiment 1

#### Bodyweight

Adult exposure groups weighted significantly more than adolescent exposure groups throughout the experiment (Table 1), including during the first (F_1,36_ = 3230.5, p < .001) and last day of drug treatment (F_1,36_ = 1083.6, p < .001), and when rats began behavioral testing (F_1,36_ = 14.7, p < .001). Bodyweights were balanced across drug groups on the first day of treatment (Drug: F_1,36_ =.91, p = .35; Drug x Age: F_1,35_ = .04, p = .85). THC exposure temporarily reduced bodyweight in both age groups, an effect that was observed on the last day of drug treatment (Drug: F_1,36_ = 20.31, p < .001; Drug x Age: F_1,36_ = .06, p = .81) but had dissipated by the time rats began behavioral testing (Drug: F_1,36_ = 1.89, p = .18; Drug x Age: F_1,36_ = .37, p = .55).

**Table 1.** Bodyweights (mean ± SEM) for groups in Experiments 1 and 2 on the first and last day of drug treatment and on the day prior to behavioral testing.

| Group |  | First injection | Last injection | Prior to testing |
| --- | --- | --- | --- | --- |
| Experiment 1 | Adolescent-THC | 114.2 ± 2.2 | 188.7 ± 3.4 | 401.1 ± 12.1 |
| | Adolescent-Vehicle | 119.3 ± 3.8 | 216.7 ± 6.4 | 441.8 ± 20.1 |
| | Adult-THC | 368.6 ± 6.1 | 384.1 ± 6.8 | 481.2 ± 16.4 |
| | Adult-Vehicle | 372.0 ± 4.9 | 409.2 ± 6.4 | 493.4 ± 14.3 |
| Experiment 2 | Adolescent-THC | 130.8 ± 6.1 | 189.1 ± 4.5 | 427.8 ± 9.4 |
| | Adolescent-Vehicle | 135.0 ± 8.5 | 204.6 ± 3.1 | 428.2 ± 10.7 |
| | Adult-THC | 400.7 ± 6.1 | 403.9 ± 7.2 | 500.2 ± 13.9 |
| | Adult-Vehicle | 399.2 ± 10.3 | 425.8 ± 12.6 | 504.8 ± 13.7 |

#### Initial action-outcome training

Rats were food deprived and trained to perform two lever-press actions for distinct food outcomes (Figure 1A), which were delivered according to an RR schedule that increased over days, such that task performance became progressively more effortful. As can be seen in Figure 1B, rats adjusted to the increase in effort by increasing their rate of lever pressing (main effect of schedule: F_3,108_ = 115.17, p < .001). Response rates were significantly elevated for THC exposed rats (drug: F_1,36_ = 5.22, p = .03), regardless of exposure age (drug x age interaction: F_1,36_ = .74, p = .40). No other effect or interaction reached significance (largest F value F_3,108_ = 1.86, p = .14 for the schedule x drug interaction).

**Fig. 1.**
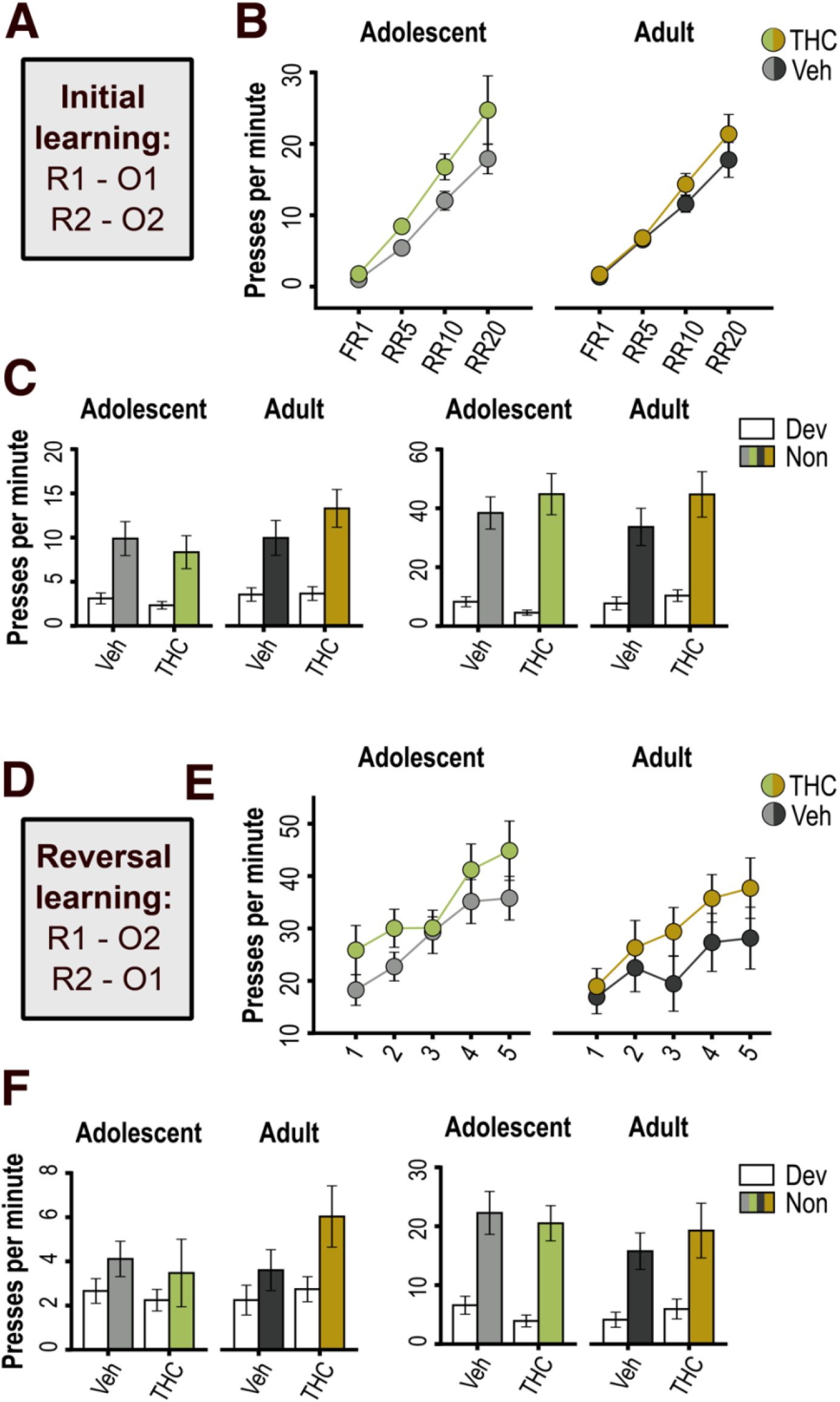
Acquisition and goal-directed control of instrumental reward seeking behavior. (A) Instrumental Response-Outcome contingencies used during initial training. (B) Response rates during initial training plotted as an average for each reinforcement schedule. (C) Response rates during the extinction (left) and reinforced (right) phases of the initial round of reward devaluation testing, plotted separately for the actions associated with the devalued (Dev) and nondevalued (Non) outcomes, as indicated. (D) Instrumental contingencies used during reversal training. (E) Response rates during reversal training sessions. (F) Response rates during the second round (post-reversal) of reward devaluation testing, as above. Data are shown as mean ± SEM for groups previously exposed to THC or vehicle (Veh) as adolescents or adults, as indicated. FR = fixed ratio, RR = random ratio.

#### Reward devaluation test

Rats then underwent reward devaluation testing to determine whether THC exposure altered their ability to use Action-Outcome associations for goal-directed decision making. As shown in Figure 1C (left panel), rats selectively suppressed their performance of whichever action earned the devalued reward during the extinction phase of the test (F_1,36_ = 57.97, p < .001), demonstrating their capacity for flexible goal-directed action selection. No effects of THC exposure (Drug: F_1,36_ = .06, p = .80; Drug x Devaluation: F_1,36_ = .43, p = .52) or Age (Age: F_1,36_ = 2.27; p = .14; Age x Devaluation: F_1,36_ = .75, p = .39) were detected, nor did these factors significantly interact (Drug x Age: F_1,36_ = .75, p = .39; Drug x Age x Devaluation: F_1,36_ = 1.11, p = .30). Sensitivity to reward devaluation was also apparent during the reinforced test phase (F_1,36_ = 104.86, p < .001), with no other significant effects or interactions (largest F value = 2.10, p = .16 for Devaluation x Drug interaction).

#### Action-outcome reversal training and reward devaluation

We next assessed the effects of THC exposure on rats’ ability to remap Action-Outcome associations during reversal training (Figure 1D) and use these updated associations to select actions based on expected reward value. Press rates (Figure 1E) increased over reversal training sessions (Session: F_4,144_ = 29.32, p < .001) and were marginally elevated in THC pretreated rats (Drug: F_1,36_ = 2.83, p = .10). No other effects were apparent (largest F value = 1.26, p = .29 for the Session x Drug x Age interaction).

As shown in Figure 1F, rats used recent Action-Outcome mappings to select actions during the reward devaluation test. For the extinction phase, there was a significant effect of Devaluation (F_1,36_ = 8.98, p = .005) and no effect of Age (F_1,36_ = .55, p = .46) or THC exposure (F_1,36_ = .42, p = .52), nor were there any significant interactions involving these factors (largest F value = 1.92, p = .17 for Age x Drug interaction). Press rates during the reinforced test phase were also sensitive to reward devaluation (F_1,36_ = 44.77, p < .001), with no other effects or interactions reaching significance (largest F value = 1.48, p = .23 for the Age x Drug interaction).

#### Contingency degradation training and testing

We then assessed the effects of THC on rats’ sensitivity to Action-Outcome contingency degradation. Rats were first retrained with the original Action-Outcome contingencies to reestablish these associations. Press rates (Figure 2A) tended to increase over days (F_1,72_ = 33.27, p < .001). THC exposed rats continued to exhibit significantly higher response rates (Drug: F_1,36_ = 4.35, p = .04) regardless of age of exposure (Drug x Age interaction: F_1,36_ = 0.00, p = .99). Apart from a marginally significant effect of age (F_1,36_ = 3.02, p = .09), no other effects or interactions were significant (largest F value = 2.06, p = 0.14 for the Day x Drug interaction).

**Fig. 2.**
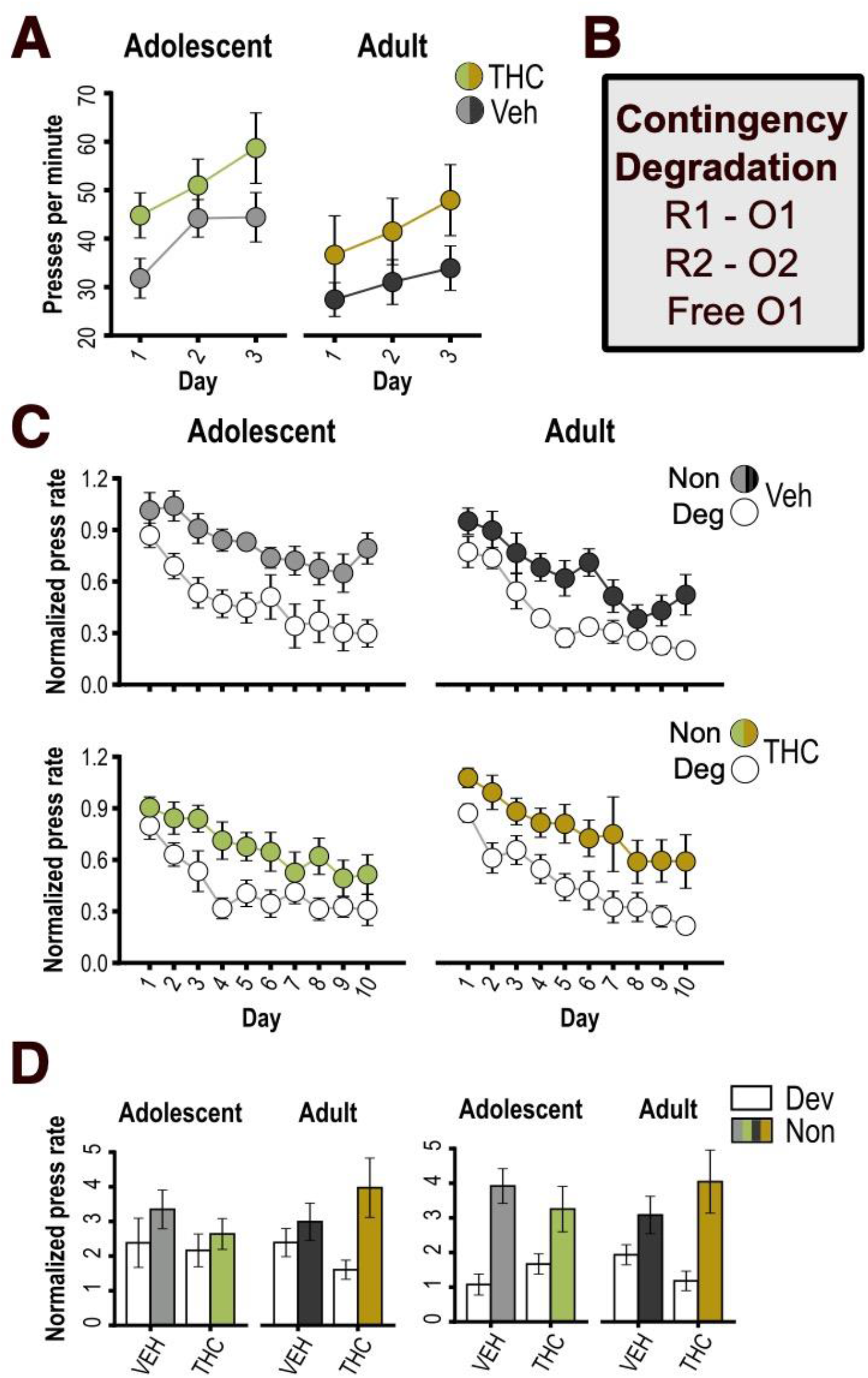
Instrumental retraining and instrumental contingency degradation learning. (A) Response rates during retraining sessions after reward devaluation testing. (B) Instrumental Response-Outcome contingencies used during instrumental contingency degradation training, indicating that one of the two outcomes was delivered in both a response-contingent and noncontingent manner. (C) Baseline-normalized response rates during instrumental contingency degradation training sessions, plotted separately for the actions associated with the degraded (Deg) and nondegraded (Non) contingencies, as indicated. (D) Baseline-normalized response rates during the early (left) and late (right) contingency degradation test sessions, plotted separately for each action as above. Data are shown as mean ± SEM for groups previously exposed to THC or vehicle (Veh) as adolescents or adults, as indicated.

During subsequent contingency degradation sessions, one of the two outcomes was delivered in a response-independent manner, such that the corresponding action was no longer a reliable predictor of reward delivery (Figure 2B). Press rates were normalized to baseline levels during the pre-training phase to adjust for individual differences. As indicated in Figure 2C, all groups reduced their overall rate of responding over contingency training sessions (Session: F_9,324_ = 53.26, p = < .001; Age x Session: F_9,324_ = 1.07, p = .38; Drug x Session: F_9,324_ = .73, p = .69; Drug x Age x Session: F_9,324_ = .37, p = .95), but also showed a selective reduction in performance of the non-predictive action (Degradation: F_1,36_ = 60.76, p < .001;Age x Degradation: F_1,36_ = .05, p = .82; Drug x Degradation: F_1,36_ = .03, p .86; Age x Drug x Degradation: F_1,36_ = 1.43, p = .24; Session x Degradation: F_9,324_ = 1.87, p = .06). No other main effects or interactions reached significance, though there was a marginal Drug x Age interaction (F_1,36_ = 2.97, p = 0.09).

Two extinction tests were conducted to probe the effects of contingency training on choice behavior in the absence of reward delivery (Figure 2D). During the first test, which was conducted midway through training, there was a general preference for the predictive (i.e., nondegraded) action (Degradation: F_1,36_ = 6.13, p = .02), with no significant differences across groups (Degradation x Age x Drug: F_1,36_ = 1.61, p = .21). No other effects or interactions were significant (largest F = 0.74, p = .40 for Degradation x Age interaction). During the final test, while there was an overall preference for the predictive action (Degradation: F_1,35_ = 35.15, p < .001), the strength of this effect was affected by THC exposure in an age-dependent manner (Degradation x Age x Drug: F_1,35_ = 4.36, p = .04; F < 1 for all other effects and interactions). This appeared to be driven by a facilitation of contingency learning in the Adult-THC group relative to their age-matched control group. Specifically, when data from the vehicle groups were analyzed separately, we found a significant Age x Degradation interaction (F_1,18_ = 4.50, p < .05). While the adolescent-vehicle group displayed a strong preference for the predictive action (F_1,9_ = 40.20, p < .001), the adult-vehicle group had yet to develop such a preference by the final test (F_1,9_ = 2.99, p = .12), suggesting a delay in contingency learning. In contrast, THC-exposed rats showed a significant preference for the predictive action (F_1,17_ = 13.88, p = .002) that did not interact with age (F_1,17_ = 1.14, p = .30). Thus, the adult-THC developed a strong preference for the predictive action (F_1,8_ = 11.58, p = .009) under conditions that did not support such an effect in the age-matched vehicle control group, suggesting an augmentation of contingency learning.

### Experiment 2

#### Bodyweight

Adult exposure groups weighed significantly more than adolescent exposure groups throughout the experiment (Table 1), including on the first (F_1,35_ = 1164.8, p < .001) and last day of drug treatment (F_1,35_ = 856.8, p < .001), as well as when rats began behavioral testing (F_1,35_ = 38.4, p < .001). Drug groups did not differ in bodyweight on the first day of treatment (Drug: F_1,35_ = .03, p = .86; Drug x Age: F_1,35_ = .13, p = .72). THC once again temporarily reduced bodyweight in both age groups by the last day of treatment (Drug: F_1,35_ = 6.30, p = .02; Drug x Age: F_1,35_ = .18, p = .67), with no such effect apparent by the beginning behavioral testing (Drug: F_1,35_ = .04, p = .84; Drug x Age: F_1,35_ = .03, p = .86).

#### Hedonic feeding

We first assessed the long-term effects of THC pre-exposure on consumption of a palatable SCM solution. Figure 3A shows the time course of SCM licking behavior (averaged across test sessions), which exhibited a typical within-session satiety profile, with reward intake peaking early in the session before entering a phase of steady decline (main effect of Time: F_29,1015_ = 67.48, p < .001). Licking behavior was not affected by THC exposure (Drug: F_1,35_ = .16, p = .70; Drug x Age: F_1,35_ = .59, p = .45; Drug x Time: F_29,1015_ = .46, p = .99; Drug x Age x Time: F_29,1015_ = 1.06, p = .38). While age groups showed similar levels of total (bodyweight-normalized) intake (F_1,35_ = .05, p = .82), the time course of consumption did significantly interact with age (F_29,1015_ = 3.55, p < .001), with older rats showing a more rapid peak and sharper decline during the early phase of intake.

**Fig. 3.**
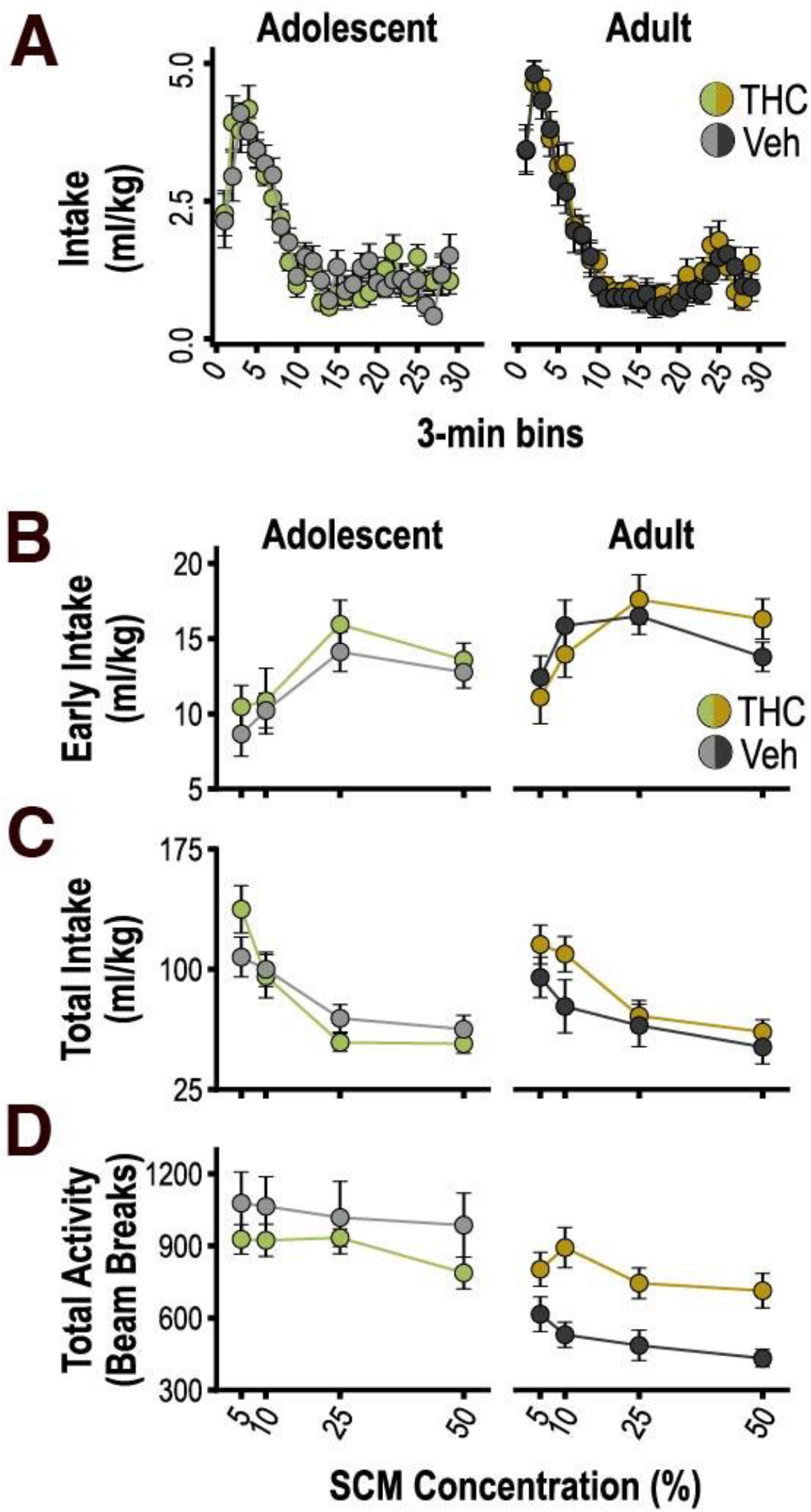
Hedonic feeding behavior. (A) Bodyweight-normalized intake of sweetened condensed milk (SCM) over time (3-min bins) during 90-min free-feeding sessions, averaged across tests. (B) Bodyweight-normalized intake during the first 3-min of active feeding, prior to satiety induction, plotted according to SCM concentration. (C) Total bodyweight-normalized intake plotted according to SCM concentration. (D) Total beam breaks during feeding sessions plotted according to SCM concentration. Data are shown as mean ± SEM for groups previously exposed to THC or vehicle (Veh) as adolescents or adults, as indicated.

To selectively assay hedonically controlled feeding and control for variability in the initiation of SCM consumption, we calculated intake during the first 3 min of active licking (i.e., after initial contact and before satiety induction; Figure 3B). As expected, early intake increased with SCM concentration (Concentration: F_3,105_ = 12.82, p < .001), reflecting the influence of palatability.

Importantly, THC pre-exposure did not alter this measure of hedonic feeding in either age group (Drug: F_1,35_ = .49, p = .49; Drug x Concentration: F_3,105_ = .73, p =.54; Drug X Age: F_1,35_ = .35, p = .56; Drug x Age x Concentration: F_3,105_ = .74, p = .53), though, as noted above, the adult exposure groups displayed generally higher levels of early intake (Age: F_1,35_ = 7.05, p = 0.01).

Total intake (Figure 3C) during consumption sessions was negatively related to SCM concentration (Concentration: F_3,105_ = 72.84, p < .001), consistent with more concentrated (and calorie-dense) solutions producing greater satiety. This measure was not significantly altered by THC exposure in either age group (Drug: F_1,35_ = .45, p = .51; Drug x Age: F_1,35_ = .17, p = .69; Drug x Concentration: F_3,105_ = .81, p = .49; Drug x Age x Concentration: F_3,105_ = .49, p = .69). Nor were there differences between age groups (Age: F_1,35_ = .17, p = .69; Age x Concentration: F_3,105_ = .58, p = .63). Thus, we found no evidence of a long-term impact of THC pre-exposure on hedonic feeding behavior or its modulation by satiety.

However, THC exposure did alter locomotor activity at test, as shown in Figure 3D. As with total intake, locomotor activity decreased as a function of SCM concentration (Concentration: F_3,105_ = 6.04, p < .001), suggesting a relationship to feeding and satiety. There was a nonspecific effect of age on locomotor behavior (Age: F_1,35_ = 16.16, p < .001; Age x Concentration: F_3,105_ = .14, p = .67), with adolescent exposure groups showing higher levels of activity. Moreover, there was an age-specific effect of THC exposure on locomotor activity (Drug: F_1,35_ = .69, p = .41; Drug x Age = F_1,35_ = 7.15, p = .01; Drug x Concentration F_3,105_ = .76, p = .52; Drug x Age x Concentration: F_3,105_ = .84, p = .48). Specifically, rats exposed to THC as adults showed higher levels of locomotor activity than the adult-vehicle group (Drug: F_1,17_ = 12.90, p = .002), whereas no such effect was observed following adolescent THC exposure (Drug: F_1,18_ = 1.16, p = .30).

#### Progressive ratio

We then assessed the effect of THC preexposure on rats’ motivation to work for SCM reward on an instrumental PR task. Once stable PR performance was established, rats were given separate PR tests with 5% or 50% SCM reward (Figure 4A). As expected, PR performance was strongly influenced by SCM concentration, with rats earning significantly more 50% than 5% reward (Concentration: F_1,35_ = 35.40, p < .001). Although PR performance appeared to be elevated in THC pretreated rats, particularly in the adult exposure condition, our analysis did not detect a significant effect of Drug (F_1,35_ = 2.13, p = .15) or Age (F_1,35_ = .16, p = .67), nor were there any significant interactions involving these factors (Age x Drug: F_1,35_ = 1.88, p = .18; Concentration x Drug: F_1,35_ = 1.49, p = .23; Concentration x Age: F_1,35_ = .06, p = .81; Concentration x Drug x Age: F_1,35_ = .06, p = .81). Given the trends in the data, Bonferroni-corrected post-hoc tests were conducted to assess the effect of THC for each age group (significance set at 0.025 = 0.05/2). The effect of THC was marginal in the adult-exposure condition (p = .04) and well above the threshold for significance in the adolescent-exposure condition (p = .95).

**Fig. 4.**
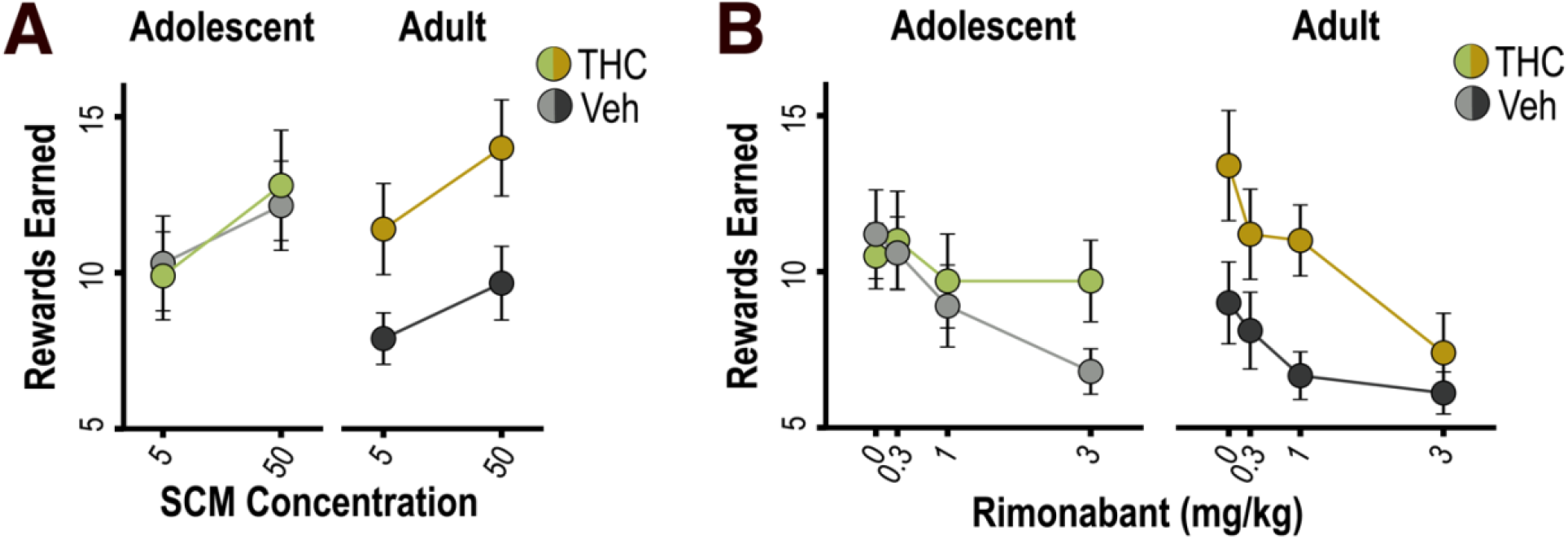
Progressive ratio testing. (A) Number of rewards earned during 90-min progressive ratio (PR) tests plotted according to the concentration of SCM used as the reinforcer. (B) Effect of rimonabant pretreatment on PR performance (total rewards earned) plotted across tests as a function of drug dose. Data are shown as mean ± SEM for groups previously exposed to THC or vehicle (Veh) as adolescents or adults, as indicated.

#### Effects of rimonabant on progressive ratio performance

We administered an additional series of tests to further probe the motivational effects of THC pre-exposure and its neurochemical basis. THC exposure is known to cause long-term changes in the expression and function of the CB1 receptor (Kruse et al. 2019; Rubino et al. 2015; Stringfield and Torregrossa 2021b; Zamberletti et al. 2012), which plays an important role in regulating reward-motivated PR performance (Friemel et al. 2014; Hernandez and Cheer 2012; Maccioni et al. 2008; Marusich and Wiley 2012; Rasmussen and Huskinson 2008; Solinas and Goldberg 2005; Ward and Dykstra 2005). We hypothesized that THC exposure might therefore alter the CB1 receptor-dependence of motivated behavior. To test this possibility, rats were pretreated with varying doses of rimonabant (0, .3, 1, 3 mg/kg) prior to PR testing. An orthogonal polynomial trend analysis was conducted to characterize linear and nonlinear dose-response functions (Randall et al. 2011; Wickens and Keppel 2004).

Rimonabant resulted in a dose-dependent reduction in the number of rewards earned on the PR task (Fig 4B), as indicated by a significant linear trend (F_1,35_ = 22.59, p < .001) and a marginally significant quadratic trend (F_1,35_ = 3.93, p = .06) of dose. There were also marginally significant linear (F_1,35_ = 3.30, p = .08) and quadratic Dose x Drug x Age interactions (F_1,35_ = 3.41, p = .07), suggesting that the groups differed in their sensitivity to rimonabant. No other effects or interactions were detected (p’s > .25), apart from a marginally significant main effect of THC exposure (F_1,35_ = 3.8, p = .06), which was related to a THC-induced elevation in PR performance in the adult-exposure condition (F_1,17_ = 5.40, p = .03) but not in the adolescent-exposure condition (F_1,18_ = .29, p = .60). When trend analyses were conducted separately for each group, both adult- and adolescent-vehicle groups exhibited a linear trend of rimonabant dose (adult: F_1,8_ = 8.30, p = .02; adolescent: F_1,9_ = 11.06, p = .009), indicating a steady decline in PR performance with increasing drug doses. In contrast, a significant quadratic trend (F_1,8_ = 9.42, p = .01) was detected for the adult-THC group, suggesting greater sensitivity to rimonabant, characterized by a sharper decline in PR performance. The adolescent-THC group also differed from controls, in this case showing diminished sensitivity to rimonabant (linear trend: F_1,9_ = 2.32, p = .16; quadratic trend: F_1,19_ = .17, p = .67).

## Discussion

Previous research indicates that chronic exposure to high doses of THC during adolescence can blunt reward processing (Realini et al. 2011; Rubino et al. 2008; Scherma et al. 2016). Less is known about how reward-related behavior is impacted by more moderate THC exposure. The current study investigated this issue using a two-week regimen of once-daily 5 mg/kg THC injections, which has been shown to support human-relevant levels of drug exposure (Ruiz et al. 2021; Torrens et al. 2020). We found that neither adolescent nor adult THC exposure caused long-term effects on hedonic feeding behavior or its regulation by satiety. Interestingly, both treatments tended to increase, rather than decrease, the vigor of instrumental performance for palatable food reward. This latter finding is consistent with recent reports that cue-motivated behavior is elevated in adulthood following adolescent exposure to relatively low doses of THC (Kruse et al. 2019; Orihuel et al. 2021). Such findings suggest that heavy and more moderate THC dosing regimens may have distinct long-term effects on reward processing and motivated behavior.

Our findings are generally in line with the view that drug-induced adaptations in the brain’s motivational hardware, particularly within the mesolimbic dopamine system, can lead to a long-term uptick in the desire to pursue rewards (i.e., ‘wanting’)(Berridge and Robinson 2016; Robinson and Berridge 1993). This increased motivation is thought to drive compulsive drug-seeking behavior but may also spillover to increase the pursuit of other nondrug rewards. For instance, previous studies have shown that rats with a history of cocaine, amphetamine, or morphine exposure exhibit an increased willingness to exert effort for palatable food reward (Forouzan et al. 2021; Mendez et al. 2009; Nordquist et al. 2007b; Rouibi and Contarino 2012), as was observed in the current study after repeated THC exposure. This heightened motivation is thought to be mediated by increased mesolimbic dopamine signaling (Berridge and Robinson 2016; Robinson and Berridge 1993), which is consistent with previous reports that repeated THC exposure causes long-lasting hyperactivity in mesolimbic dopamine neurons (De Felice and Laviolette 2021; Renard et al. 2017a), and sensitizes dopamine release in the nucleus accumbens core (Cadoni et al. 2008).

Nonspecific motivational changes arising from drug exposure may reflect an aberrant incentive process that can increase vulnerability to substance use disorder or other pathological forms of reward seeking. While there have been numerous reports of cannabinoid pre-exposure increasing voluntary opioid intake (Biscaia et al. 2008; Ellgren et al. 2007; Norwood et al. 2003; Solinas et al. 2004; Spano et al. 2007; Tomasiewicz et al. 2012; Vela et al. 1998; Ferland et al. 2022), this effect does not appear to increase willingness to exert effort for opioid reward on a progressive ratio schedule (Biscaia et al. 2008; González et al. 2004; Solinas et al. 2004). However, such studies have explored a limited range of drug pre-exposure regimens mostly involving potent synthetic cannabinoids or high doses of THC (but see Ferland et al. 2022; González et al. 2004), which should encourage further research on this topic.

Our findings suggest that the long-term effects of THC on motivation are at least partly influenced by exposure age. Progressive ratio performance was reliably elevated following adult-but not adolescent-THC exposure. These treatments also had opposing effects on the CB1 receptor dependence of progressive ratio performance, as discussed below. Furthermore, locomotor activity during consumption test sessions was elevated in the adult-THC group but not in the adolescent-THC group, even though SCM intake was not itself significantly altered by THC exposure. Locomotor activity during these sessions varied with SCM concentration and likely relates to generalized reward seeking behavior, which leads us to suggest that the elevated locomotor activity displayed by adult-THC exposed rats reflects a state of heightened behavioral arousal. These results suggest that the adolescent brain may be resilient to this long term consequence of THC exposure, which is in keeping with the recent finding that acutely administered THC is more rapidly metabolized and results in lower drug concentrations in the brain and attenuated locomotor effects in adolescent versus adult mice (Torrens et al. 2020).

THC exposure also led to more vigorous instrumental performance for food reward in Experiment 1, but in this case the effect occurred after both adolescent and adulthood THC exposure. Presumably the behavioral testing conditions used in Experiment 2 were more effective in exposing age-related variations in this persistent motivational effect of THC exposure. In Experiment 2, rats were tested on a progressive ratio schedule without food restriction in order to probe motivation related to the hedonic-emotional properties of the food reward and not its caloric-energetic properties. In contrast, rats responded for food reward on a random ratio schedule under chronic food restriction in Experiment 1 in order to support the planned testing procedures (e.g., reward devaluation through specific satiety). The use of hunger to inflate reward value in Experiment 1 may have provided a more sensitive (albeit less selective) test of motivation, thereby unmasking a potentially more subtle motivational enhancement related to adolescent-THC exposure that was not expressed in Experiment 2. This is in line with other recent studies showing that adolescent THC exposure enhances cue-motivated behavior under chronic food restriction (Kruse et al. 2019; Orihuel et al. 2021).

The endocannabinoid system is an important modulator of reward-motivated behavior (Sallam and Borgland 2021), an influence that is mediated in part by CB1-receptor-dependent facilitation of mesolimbic dopamine release (Melis et al. 2007; Oleson et al. 2012). Importantly, exogenous cannabinoid exposure triggers long-lasting adaptations in the endocannabinoid system, including in brain regions that encompass mesolimbic dopamine circuitry (Burston et al. 2010; Perdikaris et al. 2018; Rubino et al. 2015; Sim-Selley 2003; Zamberletti et al. 2012). We hypothesized that the motivational enhancement produced by adult THC exposure in Experiment 1 may reflect a change in the influence of CB1 receptor activity on reward-motivated behavior. Consistent with this prediction, we found that the response-suppressive effect of rimonabant on progressive ratio performance was altered by THC exposure in an age-dependent manner. Specifically, rats exposed to THC as adults were more sensitive to rimonabant than age-matched controls, whereas sensitivity to rimonabant was disrupted after adolescent THC exposure. These findings suggest that adult and adolescent THC exposure exert opposing effects on endocannabinergic mechanisms of motivated behavior. Previous studies have shown that adolescent THC exposure leads to widespread downregulation and desensitization of the CB1 receptor and decreases availability of its endogenous ligands anandamide and 2-arachidonoyl-sn-glycerol (Rubino et al. 2015; Zamberletti et al. 2012).

Interestingly, adolescent exposure to a relatively low-dose THC regimen (1-5 mg/kg day) is reported to selectively downregulate CB1 receptors on glutamatergic but not GABAergic synaptic terminals in the ventral tegmental area (Kruse et al. 2019). Such an adaptation would tend to disinhibit mesolimbic dopamine neurons, potentially increasing their responsivity to reinforcing and motivating stimuli. While less is known about the impact of adult THC exposure on the endocannabinoid system, it has been shown to increase CB1 receptor mRNA expression in the striatum (Romero et al. 1997), which may relate to the increased sensitivity to rimonabant observed here after adult THC exposure.

Chronic cannabinoid exposure can cause long-term deficits in cognition and higher-order executive function (Murphy et al. 2017; O’Shea et al. 2004; Quinn et al. 2008a; Schneider and Koch 2003). Goal-directed decision making is a hallmark of executive control that is essential for adaptive behavior (Balleine and O’Doherty 2010). The loss of goal-directed control may contribute to the inflexible, compulsive reward seeking that characterizes substance use disorders (Belin et al. 2013; Everitt and Robbins 2005; Ostlund and Balleine 2008). The effects of cannabinoid exposure on goal-directed behavior are not well understood. Mice given repeated THC exposure as adults have been reported to display insensitivity to reward devaluation under conditions that support flexible, goal-directed behavior in controls (Nazzaro et al. 2012). In other more recent studies, rats with a history of adolescent cannabinoid exposure displayed normal or even enhanced sensitivity to reward devaluation (Ferland et al. 2022; Orihuel et al. 2021). Importantly, these studies were focused on the transition from goal-directed to habitual control and therefore employed simple instrumental testing protocols that promote habit formation. When this approach is used, insensitivity to reward devaluation may reflect an increase in habit formation rather than a deficit in goal-directed decision making. The current study used an alternative approach in which rats were trained on a more complex two-contingency instrumental task that prevents habit formation (Colwill and Rescorla 1985; Colwill and Triola 2002; Halbout et al. 2016; Kosaki and Dickinson 2010), allowing us to more selectively assay goal-directed decision making. With this approach, we found that repeated THC exposure had no lasting effect on rats’ ability to flexibly choose between actions based on expected outcome value. This was true regardless of whether THC was administered during adolescence or adulthood, or whether testing was conducted after initial action-outcome training or after action-outcome reversal learning, which engages distinct cortical networks (Bradfield and Hart 2020; Fresno et al. 2019; Panayi and Killcross 2018; Parkes et al. 2018).

The current study also investigated the effects of THC exposure on instrumental contingency degradation training, which involves learning to withhold a goal-directed action that has lost its predictive value, and thus its utility for obtaining reward. We found that contingency degradation learning was facilitated in rats with a history of adult, but not adolescent, THC exposure. Interestingly, previous studies have found that repeated exposure to cocaine (Halbout et al. 2016) or amphetamine (Phillips and Vugler 2011) similarly enhances instrumental contingency degradation learning, suggesting that this may be a common consequence of repeated drug exposure. Future studies should probe the role of the mesocorticolimbic dopamine system in this phenomenon given that it is both crucially involved in instrumental contingency learning (Naneix et al. 2009) and persistently dysregulated by THC (Higuera-Matas et al. 2015; Perdikaris et al. 2018; Poulia et al. 2021; Renard et al. 2017a; Renard et al. 2017b; Zamberletti et al. 2012) and other abused drugs, including cocaine and amphetamine (Berridge and Robinson 2016; Pierce and Kalivas 1997; Steketee 2003).

It is notable that the enhanced contingency learning displayed by the adult-THC group reflects an improvement relative to the poor performance of the adult-vehicle group, but not relative to the otherwise good performance of the adolescent exposure groups. Unlike these other groups, the adult-vehicle group failed to selectively withhold the non-predictive (degraded) action during the final test, despite receiving ten days of contingency training. This delay in learning is not entirely surprising given that contingency degradation training was conducted after a period of reversal training, which may have created some ambiguity in the action-outcome relationships. Even when more conventional procedures are used, rats may require twelve or more days of training for the contingency degradation effect to emerge (Braun and Hauber 2012; Corbit and Balleine 2003; Corbit et al. 2002). In fact, previous reports of enhanced contingency degradation learning in psychostimulant-exposed rats also involved versions of this task that were difficult for control groups to learn (Halbout et al. 2016; Phillips and Vugler 2011). Such conditions may facilitate the detection of augmented contingency degradation learning after drug exposure. This finding may relate to previous reports that chronic low-dose THC exposure can promote neurogenesis (Cao et al. 2014; Jiang et al. 2005; Suliman et al. 2018) and improve cognition and memory in mature and aged, but not young, mice (Bilkei-Gorzo et al. 2017; Sarne et al. 2018). Such findings have raised the possibility that late-life cannabis use may have neuroprotective effects (Weinstein and Sznitman 2020).

In conclusion, the current findings demonstrate that repeated exposure to a moderate dose of THC can cause long-term motivational and cognitive effects in male rats that were generally more prominent following adult than adolescent exposure. These results highlight the need for further systematic examination of how age at exposure influences the impact of effects of chronic THC administration. Future studies should also explore the role of sex in these motivational and cognitive effects given the growing body of work identifying sex as an important factor regulating THC pharmacokinetics (Ruiz et al. 2021) and the long-term behavioral consequences of chronic THC exposure (Calakos et al. 2017; Cooper and Craft 2018).

## Acknowledgments

We thank Johanna Montesinos for assisting with animal pretreatment.

